# Cell autonomous requirement of imprinted XCI in extra-embryonic polar trophoblast cells

**DOI:** 10.1101/2022.11.10.515976

**Authors:** Feng Wang, Ashmita Chander, Yeonsoo Yoon, Mary C. Wallingford, Carmen Espejo-Serrano, Francisco Bustos, Greg M. Findlay, Jesse Mager, Ingolf Bach

## Abstract

In female mice the gene dosage from X chromosomes is adjusted by a process called X chromosome inactivation (XCI) that occurs in two steps. An imprinted form of XCI (iXCI) silencing the paternally inherited X chromosome (Xp) is initiated at the 2-4 cell stages. As extraembryonic cells including trophoblasts keep the Xp silenced, epiblast cells that give rise to the embryo proper reactivate the Xp and undergo a random form of XCI (rXCI) during peri-implantation stages. Lack of X dosage compensation leads to peri-implantation lethality due to inhibition of trophoblast stem cells. However, as the epiblast regulates the trophoblast lineage, the roles of iXCI vs rXCI in the early lethal phenotype remains unclear. We have investigated functions and expression of Rlim (Rnf12), an E3 ubiquitin ligase, and its target protein Rex1 (Zfp42) that control iXCI. Consistent with functions specifically for iXCI, we show an inverse correlation in the expression of Rlim and Rex1 throughout pre-implantation development, but an *Rlim*-independent downregulation of Rex1 in epiblast cells upon implantation. Moreover, disturbing the functional Rlim-Rex1 dynamics in females leads to cell fate confusion and premature differentiation specifically of the polar trophoblast stem cell pool. Thus, controlled by the Rlim-Rex1 axis, female mouse development requires iXCI in the polar trophoblast cell lineage.

## Introduction

Eutherian female mammals adjust the gene dosage from X chromosomes between sexes by a process known as X chromosome inactivation (XCI), a paradigm for the study of epigenetic gene silencing. In mice, XCI occurs in two waves. Beginning at the late two cell-/early four cell stage, imprinted XCI (iXCI) exclusively silences the paternal X (Xp). While extraembryonic cell types including trophoblasts maintain this XCI pattern, cells in the inner cell mass (ICM) specifically reactivate the Xp (XCR) and undergo a random form of XCI (rXCI) around implantation, where each cell inactivates either the maternal X (Xm) or the Xp (Payer, 2016). The long non-coding RNA (lncRNA) *Xist* represents a key regulator of both rXCI and iXCI. *Xist* RNA paints the inactive X from which it is expressed (Loda et al., 2022), thereby triggering downstream repressive chromatin modifications including H3K27me3 that lead to X-silencing (Plath et al., 2003).

In mice, the X-linked *Rlim* (Rnf12) gene (Ostendorff et al., 2000) is crucial for iXCI (Shin et al., 2010) but dispensable for rXCI (Shin et al., 2014). This gene encodes a RING finger E3 ligase (Ostendorff et al., 2002) that localizes mostly in the nucleus, but this cellular distribution is dependent on phosphorylation of its nuclear localization signal (NLS) (Jiao et al., 2013; Bustos et al., 2020). Female pre-implantation embryos lacking *Rlim* (KO) display severe inhibition of *Xist* transcription and defective X dosage compensation (Wang et al., 2016), as well as defective placental trophoblast development and die shortly after implantation (Shin et al., 2010). Rlim exerts its activity to induce *Xist* by proteasomal targeting of Rex1, a transcriptional repressor of *Xist* (Gontan et al., 2012). Indeed, the *Rlim* KO phenotype including iXCI is rescued in Rlim/Rex1 double KO embryos (Gontan et al., 2018), emphasizing the specificity and critical function of the Rlim/Rex1 axis in controlling iXCI. However, despite these major functions, the developmental dynamics of Rlim-Rex1 expression and the precise defects caused by axis disturbance are unknown.

During pre-implantation development Cdx2-induced trophectodermal cells (TE) form the first epithelium as outside layer of the embryo (Strumpf et al., 2005). At the blastocyst stage, TEs are segregated into mural and polar TE (mTE; pTE) with pTE cells located in close proximity to the ICM. Upon implantation mTE cells quickly differentiate into trophoblast giant cells (TGCs) that form the parietal yolk sac. In contrast, pTE cells contribute to most placental trophoblast cell types and give rise to the extraembryonic ectoderm (exe) that harbors and maintains this progenitor cell type and the ectoplacental cone (epc). The pTE progenitor state requires Fgf4 (Feldman et al., 1995; Latos and Hemberger, 2016; Tanaka et al., 1998), which is synthesized by epiblast cells, and its receptor Fgfr2c (Gotoh et al., 2005) through signaling via Raf/Mek/Erk (Arman et al., 1998; Bissonauth et al., 2006; Saba-El-Leil et al., 2003), activating a transcription factor network that includes Cdx2, Esrrb, Eomes and Elf5 among others (Auman et al., 2002; Latos et al., 2015a; Latos et al., 2015b; Luo et al., 1997; Niwa et al., 2005; Russ et al., 2000; Strumpf et al., 2005). Moreover, trophoblast development requires X dosage compensation, as deletion of *Xist* in female embryos results in loss of pTEs (Mugford et al., 2012). However, because *Xist* regulates both rXCI and iXCI, the involved mechanisms and cell types remain unclear.

Analyzing the dynamics between Rlim and Rex 1 expression, we find an inverse correlation of both proteins during preimplantation development that is severed upon implantation specifically in the epiblast cell lineage. Moreover, we show that inhibition of iXCI via KO of *Rlim* results in cell fate confusion specifically of pTEs in the exe. Our data are consistent with crucial functions of the Rlim-Rex1 axis during iXCI, but not the process of rXCI, and identify an essential role of iXCI in pTE function for successful development.

## Results

### Inverse correlation of Rlim and Rex1 expression during pre-implantation development

To investigate developmental roles of the Rlim-Rex1 axis in regulating iXCI, we first examined the expression profiles of both proteins during pre-implantation development. To obtain high quality antibodies, we developed a polyclonal Rex1 antibody raised in sheep recognizing aa 1-288 of mouse Rex1 (Segarra-Fas et al., 2022). Indeed, when tested on male embryonic stem cell (ESC) models by Western blot or by immunostaining, the Rex1 antiserum recognized Rex1 in nuclei of WT ESCs and ESCs carrying a *Rlim* deletion, but not ESCs carrying an additional Rex1 deletion (Suppl. Figs. 1A, B, respectively). The levels of Rex1 detected in single *Rlim* KO ESCs were increased when compared to WT cells, consistent with Rlim’s E3 ligase activity regulating Rex1 levels (Gontan et al., 2012). These results reveal not only a high specificity of the Rex1 antiserum but also illustrate that the Rlim-Rex1 axis is active in male ESCs.

To illuminate the dynamics of Rlim and Rex1 expression during iXCI *in vivo*, we performed co-immunostaining on whole embryos, using this Rex1 antibody in conjunction with a previously described rabbit antibody recognizing the N-terminal portion of Rlim (Wang et al., 2017), comparing animals lacking *Rlim* with WT controls. To efficiently generate embryos lacking *Rlim*, we crossed females carrying a maternal Sox-2-Cre mediated *Rlim* cKO and a paternal *Rlim* KO allele (*Rlim* flox_m_/KO_p_; Sox2-Cre^+/-^) with males lacking *Rlim* (KO/Y; Suppl. Fig. 1C). This mating strategy ensures that all offspring lack *Rlim* (Shin et al., 2014; Wang et al., 2016). We first focused our analyses on whole pre-implantation embryos stages E2.5, E3.5 and E4.5. Because *Rlim* is among X-linked genes that are quickly silenced during the iXCI process, it is mostly expressed from the maternal allele at these stages both in female and male offspring (Borensztein et al., 2017), and thus, Rlim effects on Rex1 levels in male and female embryos are comparable. This strategy allows an interrogation of expression dynamics in tissues that are affected by iXCI in females, such as cells of the trophoblast lineage. As previously reported (Shin et al., 2010; Shin et al., 2014), in WT embryos Rlim is expressed throughout pre-implantation development (Fig. 1). Likewise, Rex1 expression was low but detectable in these embryos at all stages (Fig. 1), with strictly nuclear localization. At E2.5 Rex1 protein was also detected in specific nuclear punctae that, as judged by comparison with DAPI staining, did not correspond to centric heterochromatin domains. Rlim protein localization in cells was somewhat more diffuse indicating nuclear and some cytoplasmic localization, suggesting shuttling between these compartments (Jiao et al., 2013), co-localizing with Rex1 in the nucleus. Starting at E4.5 and thereafter, Rlim localization appeared even more diffuse specifically in epiblast tissues (Figs. 1; 2), as previously reported (Shin et al., 2014). Even though immunostaining is non-quantitative, in embryos lacking *Rlim*, we detected elevated Rex1 immunoreactivity throughout pre-implantation development when compared to control, and this was particularly apparent at E3.5 (Fig. 2). These results are consistent with both, Rlim’s functions in targeting Rex1 for degradation (Gontan et al., 2012), and a gradual upregulation of Rex1 mRNA from low levels at the 4 cell stage to high levels at blastocyst stages (Wang and Bach, 2019). Interestingly, expression of Rex1 in nuclear punctae of E2.5 embryos appeared resistant to influences of Rlim activity. Investigating expression patterns in whole embryos at early post-implantation stages, we found Rlim robustly expressed in extraembryonic tissues of WT embryos, but weak cytoplasmic Rlim still detectable in the epiblast at E5.5 but no longer at E6.5 (Fig. 2). Rex1 immunoreactivity in WT embryos was very low. Importantly, while extraembryonic tissues in the *Rlim* cKO continued to exhibit increased Rex1 levels, indicating continued roles of Rlim in the turn-over of Rex1, this was no longer the case in epiblast cells (Fig. 2). Combined, these results reveal an inverse relationship between expression of Rlim and Rex1 during early mouse development, consistent with regulation of iXCI, and provide first evidence that upon implantation Rex1 downregulation in epiblast cells occurs in an *Rlim-*independent manner.

**Figure 1:**
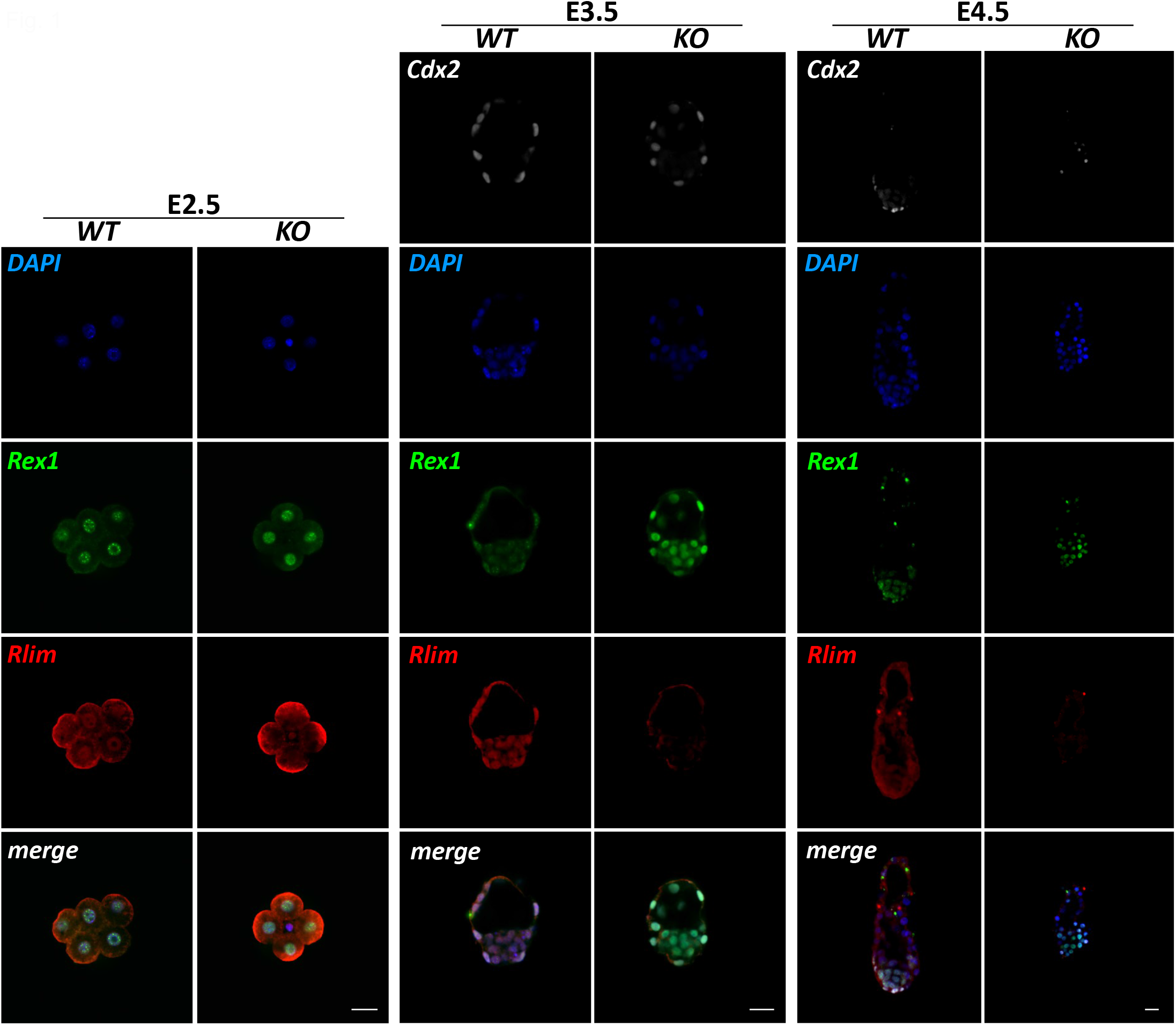
Increased Rex1 levels in pre-implantation embryos lacking Rlim. WT and *Rlim* KO embryos at stages E2.5, E3.5 and E4.5 were co-stained in parallel using indicated antibodies. Image recordings/processing of KO and control embryos were carried out with the same settings. Rlim=red; Rex1=green; Cdx2=white. DAPI=blue. Note elevated Rex1 immunoreactivity in embryos lacking *Rlim*. Scale bars=25 µm.

**Figure 2:**
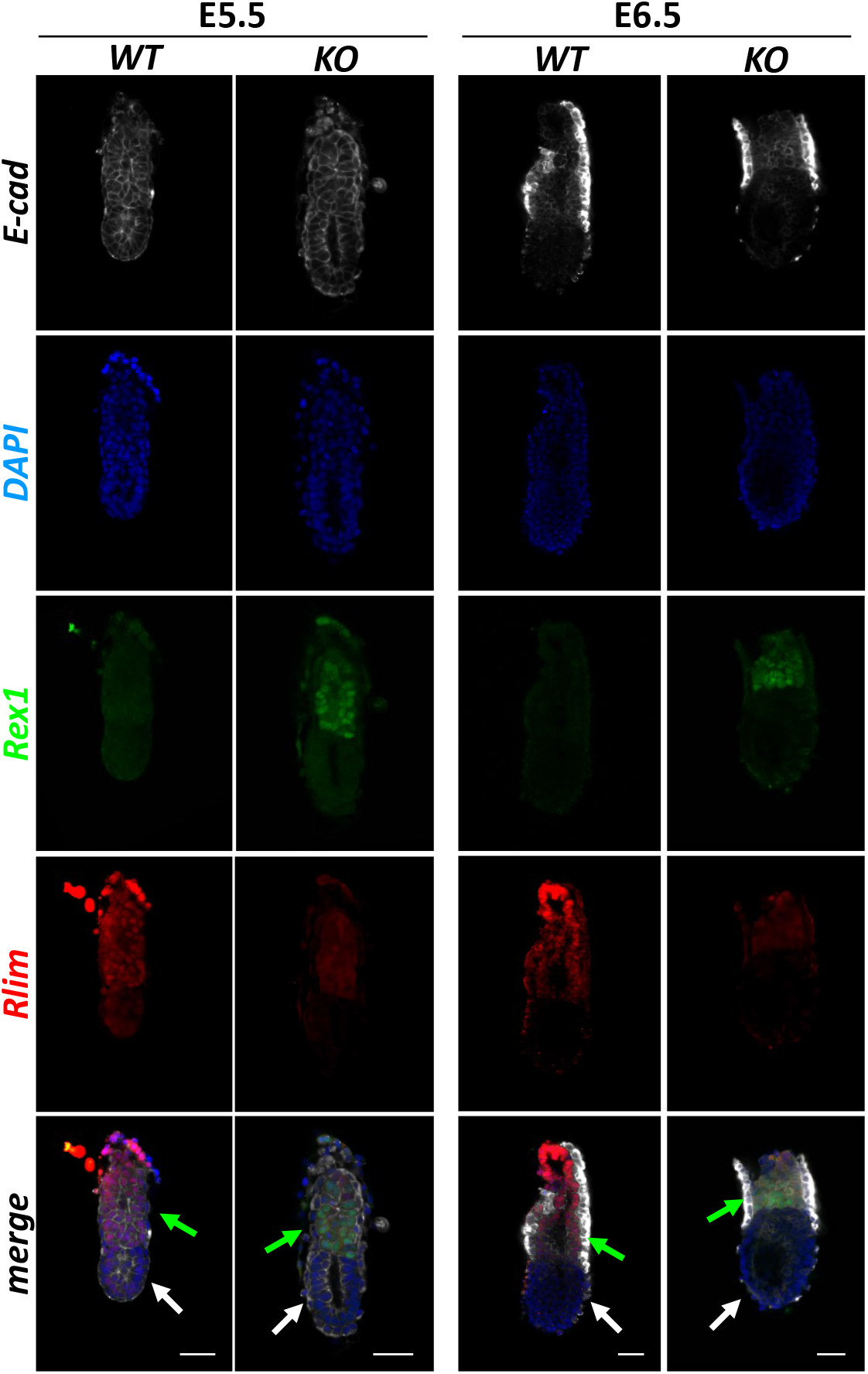
Rlim-independent downregulation of Rex1 in epiblast cells upon implantation. WT and Rlim KO embryos at post-implantation stages E5.5 and E6.5 were co-stained in parallel using indicated antibodies. Image recordings/processing of KO and control embryos were carried out using the same settings. Rlim=red; Rex1=green; Cdx2=white. DAPI=blue. Green and white arrows indicate exe and epiblast domains, respectively. Note increased Rex1 immunoreactivity specifically in trophoblast but not epiblast regions in embryos lacking *Rlim*. Scale bars =50 µm.

### iXCI is required specifically in pTE precursor cells in the exe

It is well-established that inhibition of iXCI in *Rlim* cKO females leads to lack of trophoblast tissues and peri-implantation lethality. To begin addressing the importance of iXCI, we examined the precise and sex-specific functional consequences of disturbing the Rlim-Rex1 dynamics and analyzed effects of the *Rlim* KO on extraembryonic tissues, starting with mTE cells. Previous studies suggested that lack of iXCI has no major effect on pre-implantation development (Shin et al., 2010) and thus, we first analyzed mTEs in female blastocyst outgrowths lacking *Rlim* generated by mating flox_m_/KO_p_; Sox2-Cre^+/-^ mothers with KO/Y sires (Suppl. Fig. 1C). As males do not undergo iXCI, male littermates served as controls. Blastocysts were isolated at E3.5 and cultured up to 5 days. The sex was determined via PCR, after image recording. At day 1 of culturing, mTEs stained Cdx2^+^ and no obvious differences were observed between female and male embryos (Suppl. Fig. 2). At 3 days in culture both male and female embryos had hatched, with similar numbers of mTE-derived differentiated giant cells, as previously reported (not shown)(Shin et al., 2010). Moreover, overall rates of mitosis and cell death at this stage appeared not significantly different between sexes as judged by staining with antibodies against proliferation marker phospho-histone 3 (pH3) or apoptosis marker cleaved Caspase 3, respectively (Suppl. Fig. 2; not shown). However, Cdx2 was detectable but with notably decreased signal in female embryos, suggesting defective pTE cell functions. At day 5, all male outgrowths (n=14) but none of the female outgrowths (n=12) displayed an outgrowth likely consisting of pTE-derived cells (Fig. 3). These results are consistent with published results (Shin et al., 2010; Shin et al., 2014) and suggest that major developmental effects caused by lack of *Rlim* are in pTE-derived cell types, specifically in females.

**Figure 3:**
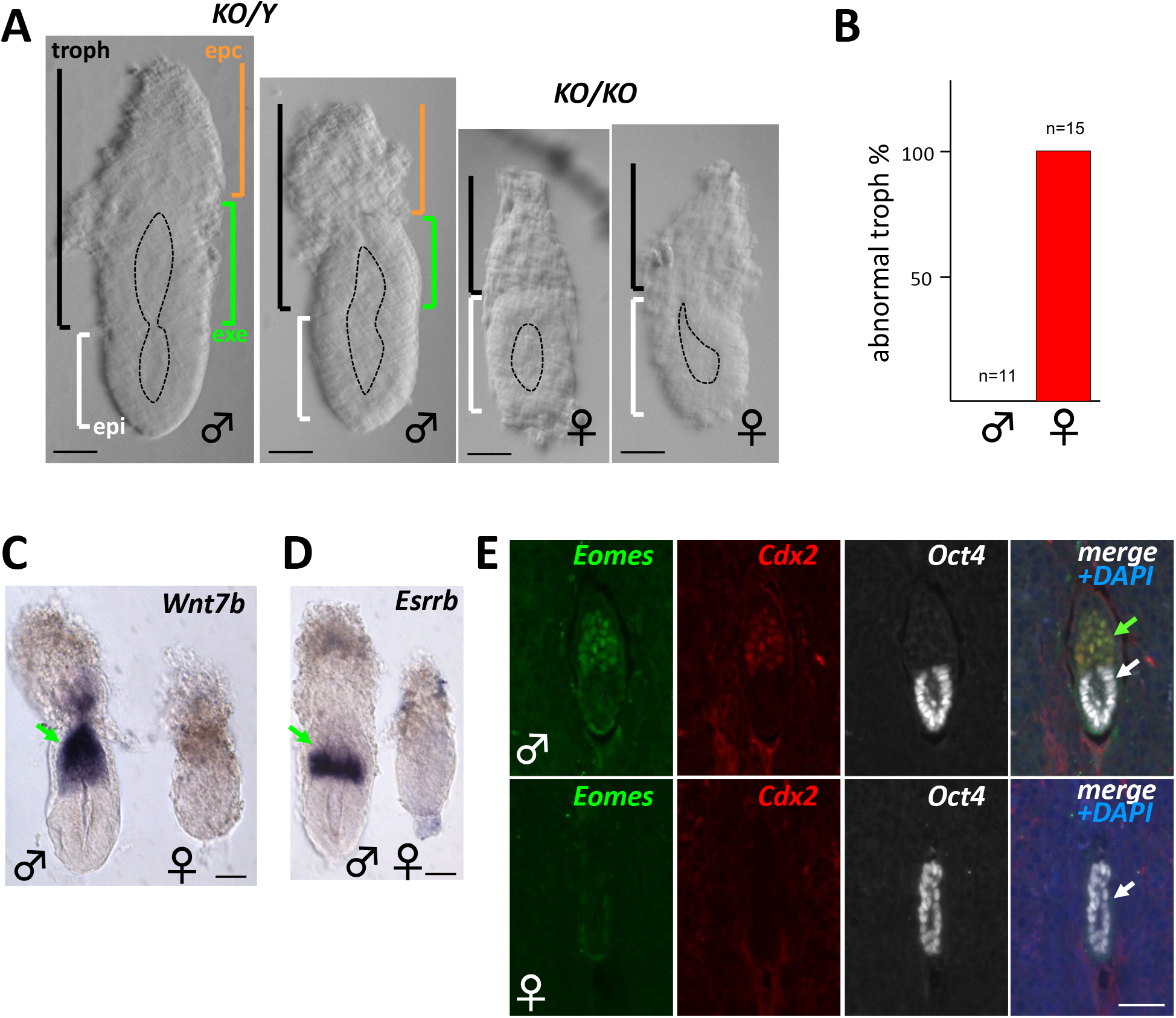
Absence of the exe structure in female embryos lacking *Rlim*. Male/female littermates within subfigures are shown at the same magnification. Gender was determined after image recording. **A)** Representative brightfield images of E5.75 embryos. Approximate embryo and lumen domains are indicated. n>27 each. epc=ectoplacental cone region (brown); exe=extraembryonic ectoderm region (green); troph= trophoblast-derived (black); epi=epiblast (white). Scale bars = 60 µm. **B)** Quantification of phenotypic appearance. **C, D)** Whole embryo *in situ* hybridization. Lack of exe markers Wnt7b (C) and Esrrb (D) in E5.75 *Rlim* KO females (n>10, each). Scale bars = 60 µm. **E)** IHC of *Rlim* KO embryonic sections at E5.5 within placentae reveals lack of exe markers Eomes and Cdx2. Genotyping after image recording. Scale bar = 40 µm. In C-E green and white arrows indicate exe and epiblast regions, respectively.

To better visualize embryonic structures including the epiblast, exe and epc, we continued our analyses on dissected early post-implantation embryos at developmental stages E5.5 and E6.5. While epiblast (epi) tissues, epiblast lumenogenesis and mTE-derived parietal yolk sac were present both in male and female embryos, presumed pTE-derived trophoblast domains appeared disorganized in 100% of females examined (Fig. 3 A, B). Using wholemount *in situ* hybridization or IF on sections of embryos within placentae failed to detect exe markers including Wnt7b (Kemp et al, 2007; Yoon et al., 2012), Esrrb (Guzman-Ayala et al., 2004; Luo et al., 1997), Eomes (Russ et al., 2000) and Cdx2 (Ralston et al., 2010; Strumpf et al., 2005)(Figs. 3C-E). The lack of these markers that are normally expressed in specific exe subdomains suggest a complete absence of pTE-derived cell types that make up the exe. In contrast, Oct4^+^ (Rosner et al., 1990; Scholer et al., 1990) epiblast tissue was present (Fig. 3E), indicating exe-specific defects in *Rlim* null female embryos.

Next, we investigated the state of XCI in female embryos lacking *Rlim*, staining sections of embryos in decidua with antibodies against H3K27me3, a marker of the inactive X chromosome (Plath et al., 2003). Because some female *Rlim* KO/KO embryos survive until E7.5 (Shin et al., 2010), which corresponds to the earliest stage when H3K27me3 foci induced by rXCI are detectable in the epiblast (Shin et al., 2014), we interrogated embryos at this stage. Indeed, female control embryos displayed H3K27me3 foci both in epiblast and trophoblast tissues (Fig. 4). Consistent with findings that *Rlim* is required for iXCI but not rXCI (Shin et al., 2014), *Rlim* KO/KO females formed H3K27me3 foci in epiblast, but not in trophoblast tissues (Fig. 5). Combined, these results provide strong evidence that the process of iXCI is specifically required for female embryos to form the exe structure.

**Figure 4:**
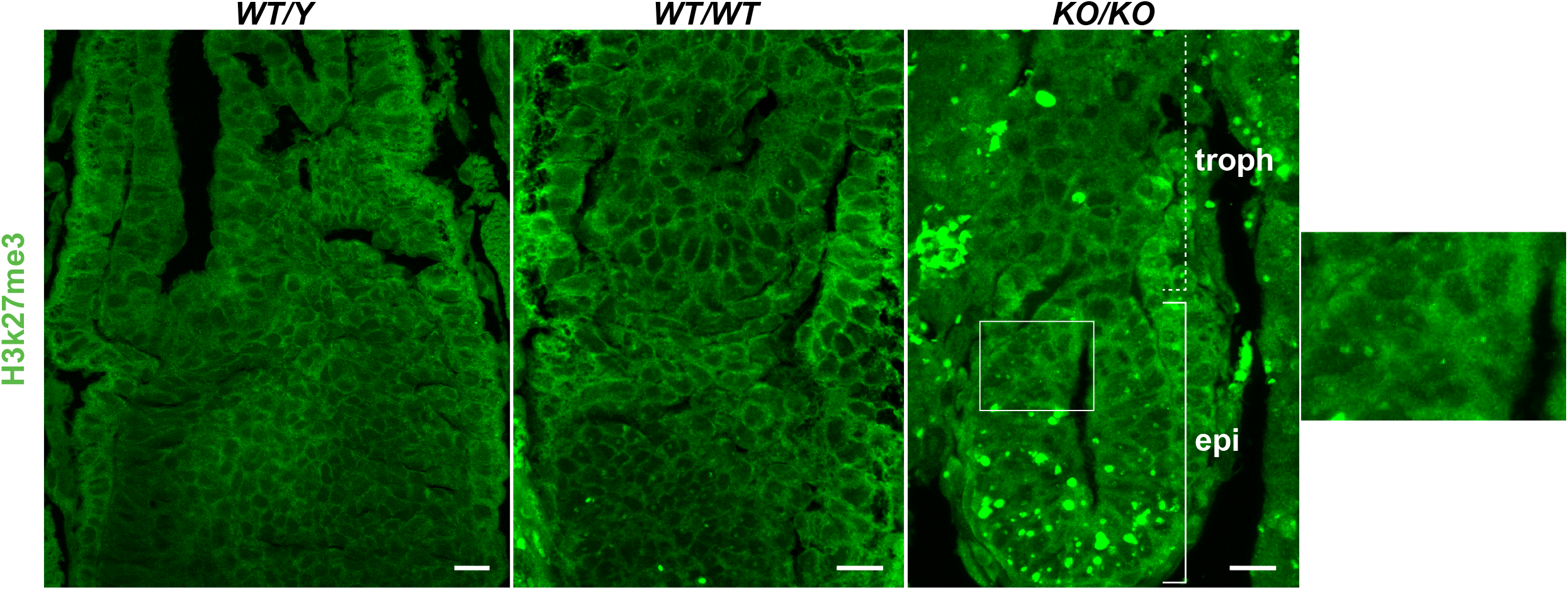
Defective iXCI in polar trophectoderm derived tissues in females lacking *Rlim*. Sections of WT/Y male, and WT/WT and *Rlim* KO/KO female embryos in placentae at stages E7.5 were stained with H3K27me3 antibodies. Shown are representative images depicting epiblast and bordering trophoblast embryonic regions. Boxed area is shown in higher magnification. Note presence of H3K27me3 foci in female KO/KO epiblast but not trophoblast tissues. Larger spots are likely caused by unspecific staining of small blood accumulations. Scale bars = 20µm.

**Figure 5:**
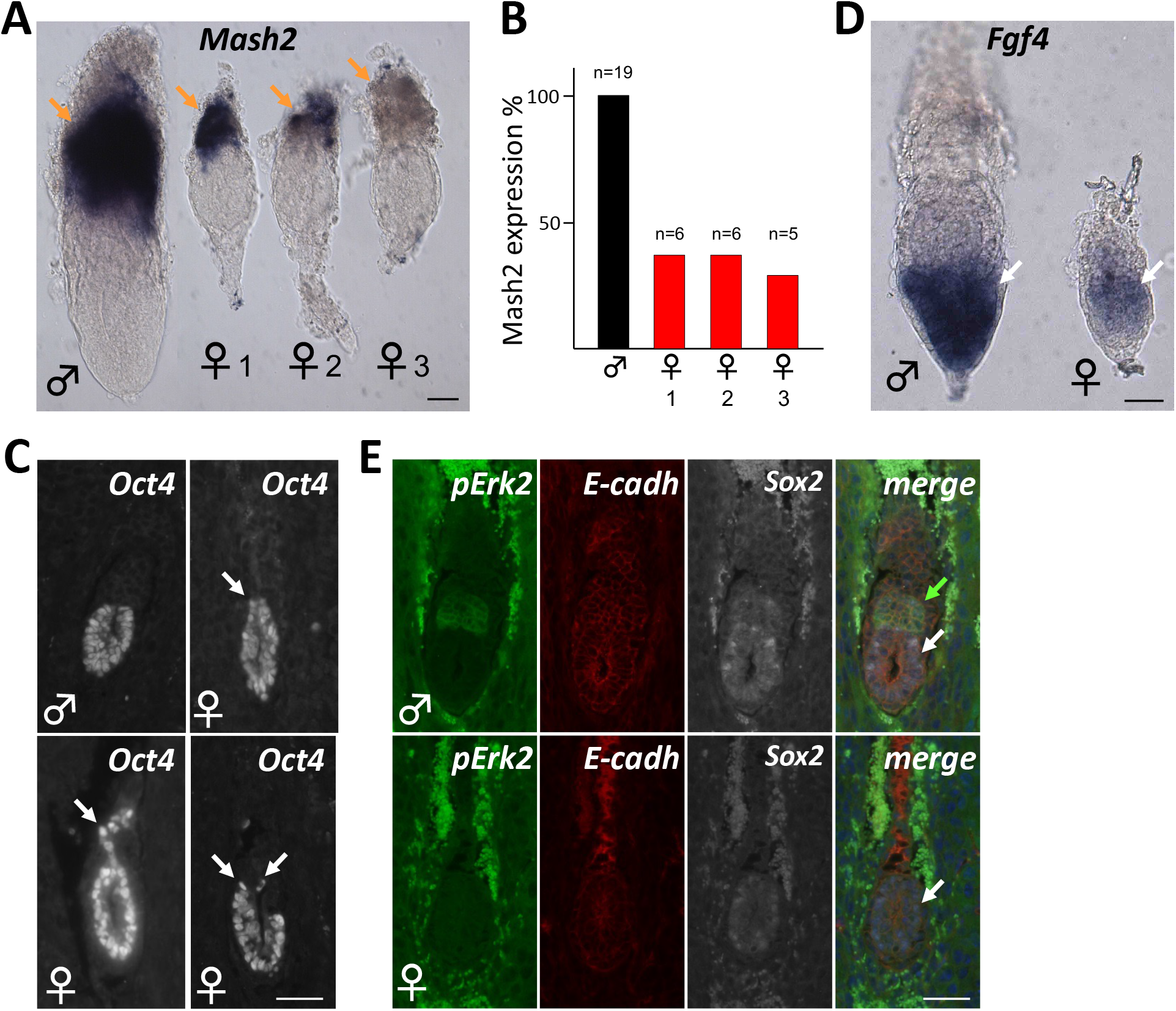
Lack of iXCI causes pTE cell fate confusion in the presence of Fgf4. Representative images of male and female *Rlim* KO littermate embryos at E6.5. A, C: whole mount ISH; B, D: IHC, sections. Genotyping after image recording. **A)** *In situ* hybridization with epc marker Mash2 reveals female embryos with strong (1), intermediate (2) and weak/no staining (3). Scale bar = 50µm. **B)** Summary of A. **C)** Examples of undefined epi/troph border and/or expansion of Oct4^+^ cells into trophoblast domains (arrows). Scale bar = 40µm. **D)** *In situ* hybridization of embryos at E6.5 using Fgf4 antisense probe. Note expression of Fgf4 in the epiblast of female *Rlim* KO embryos. Scale bar = 50µm. **E)** Absence of pErk2^+^ cells in *Rlim* KO females. Same magnification of images within sub-figures. Scale bar = 40µm. Brown arrows = epc or epc-like; white arrows = epiblast or epiblast-like; green arrow = exe.

### Lack of iXCI causes cell fate confusion of pTE precursor cells

Even though the exe structure is entirely absent in *Rlim* KO females, pTE-derived cells appear present albeit at low quantity (Fig. 3A). Examining these tissues, we did not detect elevated induction of cell death marker cleaved caspase 3 during post-implantation development (not shown). We next tested expression of Mash2 (Guillemot et al., 1994; Guzman-Ayala et al., 2004) a marker for spongiotrophoblast cell types in the epc using whole embryo *in situ* hybridization. Interestingly, while some female embryos exhibited strong staining comprising the entire trophoblast domain, partial/intermediate or no/weak Mash2 staining was detected in others (Fig. 5A), and these Mash2 expression patterns appeared in similar frequencies (Fig. 5B). We also noted that many female embryos lacking *Rlim* did not develop a defined epiblast-trophoblast boundary and that Oct4^+^ cells were detectable well into the trophoblast domain (Figs. 3E; 5C). Because Fgf4 expressed by epiblast cells is crucial for the maintenance of pTE precursor cells (Tanaka et al., 1998), we tested Fgf4 expression in female embryos via whole embryo in situ hybridization. Indeed, while the Fgf4 signal in males was more robust, female embryos at E6.5 still expressed Fgf4 mRNA, even though cells in the trophoblast compartment expressing activated phospho-Erk1,2 (p-Erk) were no longer detectable (Fig. 5D, E), indicating that loss of pTE precursors occurs in the presence of Fgf4. Combined, these results suggest that lack of dosage compensation specifically in pTE precursor cells in the extraembryonic ectoderm causes cell fate confusion and cells prematurely differentiate into various cell types including the Mash2^+^ spongiotrophoblast lineage.

## Discussion

In female pre-implantation embryos, iXCI gradually silences Xp-linked genes until X dosage compensation is reached at late blastocyst stages, including in trophoblast cells (Wang et al., 2016). To illuminate the roles of the Rlim-Rex1 axis during X dosage compensation in female mice, and its potential impact for the processes of iXCI and rXCI as well as XCR, we compared the expression of Rex1 protein in embryos with and without *Rlim*. In whole embryo staining, the overall pattern of Rex1 expression during pre-implantation in WT controls, combined with elevated Rex1 levels in *Rlim* KO animals strongly support the proposed functions of the Rlim-Rex1 axis for controlling iXCI (Gontan et al., 2018), with Rlim targeting Rex1 for proteasomal degradation (Gontan et al., 2012), thereby emphasizing the importance of the dynamics between these proteins. It is important to point out that as early as E2.5, Rex1 protein is detectable throughout pre-implantation development even when Rlim is present (Fig. 1), indicating that much of the process of iXCI occurs in the presence of Rex1. Considering that *Rex1* mRNA levels are low at early stages of mouse pre-implantation development (Wang and Bach 2019), and signs of *Xist* expression and cloud formation are present in females lacking *Rlim* by E2.5 (Wang et al, 2016), it is highly likely that Rex1 protein levels are low at the 2-4 cell stage when iXCI is initiated, even in animals lacking *Rlim*. Consistent with this, expression data from animals at E2.5 indicate low levels of Rex1, with levels only slightly increasing upon deletion of *Rlim*. In contrast, the requirement for Rlim activity to maintain Rex1 at low levels appear most critical at E3.5, when both Rex1 mRNA and *Xist* levels are particularly high (Wang and Bach, 2019; Wang et al., 2016). These results are not only fully consistent with expression data of these genes but also with the temporary X-silencing profiles as elucidated by whole embryo RNAseq (Wang et al., 2016). The presence of Rex1 protein in cells of WT animals suggests that Rlim’s E3 ligase activity is likely not only required to decrease overall nuclear levels of Rex1, but also more locally to clear Rex1 off promoters such as *Xist*, as Rlim targets DNA-bound transcription factors (Ostendorff et al., 2002). Moreover, the change of low levels of Rex1 to high levels in embryos with and without *Rlim*, respectively indicates that in the absence of *Rlim*, threshold levels of Rex1 are reached that inhibit iXCI. As females carrying a single functional *Rlim* allele (WT_m_/KO_p_) undergo normal iXCI (Shin et al., 2010), these findings provide strong evidence that it is the dose of Rex1, set by *Rlim*, that critically controls the iXCI process.

Expression data on post-implantation embryos demonstrate that the Rlim-Rex1 axis continues to be active in extraembryonic tissues, as indicated by increased levels of Rex1 in the exe of *Rlim* KO animals (Fig. 2). Because *Rlim* KO/KO females do not form this structure (Fig. 3), these results further reveal that the functional Rlim-Rex1 interactions in mice are not female-specific but active also in male embryos. Concerning the epiblast cell lineage, even though we cannot exclude a slight delay in onset, females lacking *Rlim* display signs of rXCI by E7.5, in contrast to pTE-derived extraembryonic tissues (Fig. 4). The weak and diffuse signal of Rlim at peri-implantation stages in WT embryos is consistent with cytoplasmic staining as shown previously on embryonic sections (Shin et al., 2014). Thus, upon implantation, both Rlim and Rex1 expression is specifically downregulated in nuclei of epiblast cells. This result combined with the finding that *Rlim* KO embryos do not upregulate Rex1 in the epiblast (Fig. 2), indicates that the Rlim-Rex1 axis becomes inactive between E4.5 and E5.5, when Rex1 levels in epiblast cells are downregulated in an *Rlim*-independent manner (Figs. 1, 2), thereby providing a molecular explanation of why *Rlim* is not required for rXCI (Shin et al., 2014). Moreover, our data suggests that in mice, rXCI in epiblast cells is initiated when Rex1 levels are low. As various ESC models express very different and sometimes high levels of Rex1 at the time when rXCI is induced, this resolves a controversy of the requirement/role of *Rlim* for undergoing XCI in female ESCs (Shin et al., 2010; Barakat et al., 2011; Shin et al., 2014; Wang et al., 2017). Moreover, by removing repressive functions of Rex1 on *Xist* transcription, these results may also provide an explanation of why overexpression of Rlim leads to induction of XCI in a small number of male ESCs (Jonkers et al., 2009).

Concerning an involvement of the Rlim-Rex1 axis for the XCR process, which in mice occurs between E4.5 and E5, we observed more pronounced cytoplasmic localization of Rlim in WT epiblast cells beginning at E4.5, as previously reported (Shin et al., 2014) (Fig. 2), but this did not lead to an obvious spike in levels of nuclear Rex1 (Figs. 1, 2). However, because Rlim can target DNA-bound transcription factors (Ostendorff et al., 2002), its cytoplasmic sequestering may nevertheless allow for a temporary limited, enhanced occupancy of Rex1 on *Xist* regulatory sequences and, depending on the kinetics of Rex1 downregulation, this could potentially lead to Xp reactivation. Thus, influences of the Rlim-Rex1 axes on XCR in female mice cannot be excluded.

The expression pattern obtained by staining whole embryos using an antibody recognizing the N-terminal 200aa of Rlim (N-Rlim) (Wang et al., 2017) is consistent with previous IHC using a different ab recognizing the central portion of Rlim, both at pre-implantation stages and post-implantation stages (Shin et al., 2010; 2014). However, we noticed weak but reproducible cytoplasmic signal in post-implantation embryos, specifically in extraembryonic tissues of *Rlim* cKO (Fig. 2). As the N-Rlim antibody recognizes N-terminal Rlim sequences including 83aa that correspond to a region not looped out in the cKO (Wang et al., 2017), these results suggest that a N-terminal Rlim peptide of 83aa is expressed in the cKO in some tissues. Thus, even though this peptide does not contain known functional domains, including the Rex1 interaction domain, nuclear import/export signals and the RING finger (Wang et al., 2017), and is not detectable at pre-implantation stages (Fig. 1) when iXCI occurs, we cannot fully exclude minor influences of this peptide on the *Rlim* KO phenotype in female mice.

Concerning the developmental phenotype of females lacking *Rlim*, pre-implantation development including the development of initial trophoblast and epiblast tissues appears normal, as previously reported (Shin et al., 2010; Wang et al., 2016). This indicates at least some resistance to X dosage defects, considering that iXCI affects X dosage compensation of all cell types in the female pre-implantation embryo. However, a highly penetrant lethal embryonic phenotype precipitates upon implantation, as we have not observed a single born female pup carrying a maternally transmitted *Rlim* KO allele in more than 10 years of breeding. The epiblast cell lineage does not appear to display major defects in early post-implantation embryos, as our data provides evidence that upon implantation, female epiblast tissues lacking *Rlim* undergo lumenogenesis (Fig. 3), downregulate Sox2 expression (Fig. 5), develop H3K27me3 foci (Fig. 4), and continue to express Fgf4 (Fig. 5). Moreover, differentiation of mTE-derived parietal TGCs lacking *Rlim* occurs in a similar fashion to control animals (Figs. 3; Suppl. Fig. 2), indicating limited requirement of iXCI on the differentiation and functions of these cell types. Thus, the main phenotype of the *Rlim* KO in female embryos appears restricted to the development of pTE-derived structures. The findings that extraembryonic cell types exclusively undergo iXCI combined with Fgf4 expression detected in *Rlim* KO/KO epiblast tissues (Fig. 5) provides strong evidence that a lack of X dosage compensation specifically in pTE progenitor cells is the underlying cause of the observed phenotype. The reason for such striking specificity in X dosage sensitivity by pTEs is unknown and needs further investigation. This specificity is further emphasized by the fact that pTEs appear exquisitely sensitive to the presence of *Xist* (Mugford et al., 2012), in contrast to epiblast-derived tissues (Yang et al., 2016). Thus, iXCI in pTEs may represent a major developmental bottleneck, at least in mice. As the process of iXCI is not employed in many mammalian species including humans (Okamoto et al., 2011), it will be interesting to investigate the sensitivity of pTEs for X dosage compensation across species.

Concerning the fate acquired by female pTE precursors lacking X dosage compensation, as cells occupying the trophoblast compartment were present, but we did not find evidence of cell death (data not shown)(Shin et al., 2010; Shin et al., 2014), it appears likely that these cells prematurely differentiate without self-maintenance, causing the collapse of the exe structure. The finding that pTE cells of some embryos lacking *Rlim* differentiate into Mash2^+^ cells but not those in other embryos (Fig. 5), indicates inconsistent differentiation into various cell fates, suggesting cell fate confusion, likely caused by lack of lineage-defining factors including Cdx2, Eomes and Esrrb (Latos et al., 2015a; Russ et al., 2000; Strumpf et al., 2005). Indeed, trophoblast cell lineages other than Mash2^+^ spongiotrophoblasts exist that do not express Mash2, including the labyrinthine trophoblast lineage (Tanaka et al., 1997). In this context, the extension of Oct4^+^ cells into trophoblast domains (Fig. 5C) might be caused by migration of Oct4^+^ epiblasts cells. Alternatively, considering that reciprocal repression between Oct4 and Cdx2 is important for initial trophoblast specification (Niwa et al., 2005), the downregulation of Cdx2 in female pTEs lacking X dosage compensation (Figs. 3; Suppl. Fig. 2) may lead to the reversion of Oct4 expression in some pTE cells. Thus, more work is needed to elucidate specific cell fates acquired by pTE precursor cells lacking X dosage compensation.

In summary, our data provide evidence for a decisive role of the functional dynamics between Rlim and Rex1 in the control of iXCI during female mouse embryogenesis. Moreover, our results show that defective iXCI in pTE precursor cells results in loss of cell identity causing cell fate confusion, likely leading to inhibition of self-renewal and progressive depletion of the pTE cell pool thereby abrogating implantation. Thus, cell autonomous X dosage compensation in pTEs represents a strict requirement for iXCI in pTE cells for successful mouse development.

## Material and Methods

### Mice

Mice used in this study and genotyping have been described. *Rlim fl/fl* mice and *Sox2-Cre* mice (JAX Stock #008454; in C57BL/6) as well as their genotyping have been described (Hayashi et al., 2002; Shin et al., 2010; Shin et al., 2014). *Rlim* mice were bred and maintained in a C57BL/6 background. All mice were housed in the animal facility of UMMS and utilized according to NIH guidelines and those established by the UMMS Institute of Animal Care and Usage Committee (IACUC).

### Antibodies

Concerning the generation and characterisation of the REX1 antibody, a sheep anti-mouse REX1 (1-288) antibody was raised by MRC PPU Reagents & Services (MRC PPU R&S code DA136). GST-tagged mouse REX1 (1-288) was used as the immunogen and the serum purified against MBP-tagged REX1 (1-288) to minimise purification of GST antibodies. Antibody specificity was determined by immunofluorescence and immunoblotting. For immunoblotting, *Rlim* ^+/y^, *Rlim* ^-/y^ or *Rlim* ^-/y^ ;*Zfp42* ^-/-^ mESCs (Bustos et al., 2018; Bustos et al., 2020) were cultured on gelatin and lysed in 20 mM tris-HCl (pH 7.4), 150 mM NaCl, 1 mM EDTA, 1% (v/v) NP-40, 0.5% (w/v) sodium deoxycholate, 10 mM β-glycerophosphate, 10 mM sodium pyrophosphate, 1 mM NaF, 2 mM Na_3_VO_4_, and cOmplete Protease Inhibitor Cocktail Tablets (0.1 U/ml; Roche) and protein extracts resolved by SDS-PAGE prior to transfer. REX1 antibody DA136 (3^rd^ bleed) was diluted at 1:1000 and visualized using Donkey anti-Sheep HRP secondary antibody (Thermo Fisher Scientific). Immunoblots were imaged using enhanced chemiluminescence on a BioRad ChemiDoc imager (BioRad). For immunostaining, *Rlim* ^+/y^, *Rlim* ^-/y^ or *Rlim* ^-/y^;*Zfp42* ^-/-^ mESCs were cultured on gelatin coated coverslips, fixed with 4% PFA in PBS for 20 min, permeabilized in 0.5% Triton X-100 PBS 5 min and blocked with 1% Fish gelatin PBS 30 min. REX1 antibody DA136 (3^rd^ bleed) was diluted at 1:1000 in blocking solution and coverslips were incubated for 2 h. Alexa Fluor 488 Donkey anti-Sheep IgG (Thermo Fisher Scientific) was used as secondary antibody at 1:500 in blocking solution and incubated for 1 h. DNA stain was performed with Hoechst (1:10,000 in PBS) for 5 min. Coverslips were mounted using Fluorsave reagent and imaged using a Zeiss 710 confocal microscope with Zen software (Zeiss). The N-Rlim antibody has previously been described (Ostendorff et al., 2002; Wang et al., 2017). Other commercial primary antibodies were Cdx2 (Biogenex MU392A-UC), pH3 (Millipore NG1740993), Oct4 (Abcam, 200834), E-cadherin (BD biosciences 610181), Eomes (Abcam 23345), H3K27me3 (Millipore Sigma 07-449), pErk1/2 (Cell Signaling #9102), Fgfr2 (Abcam 10648), Casp3 (Cell Signaling #9661), Sox2 (Abcam 97959). Secondary antibodies used include Alexa Fluor® 488 Donkey Anti-Rabbit IgG (Invitrogen, A21206), Alexa Fluor® 488 Goat Anti-mouse IgG (Invitrogen, A11029), Alexa Fluor® 568 Goat Anti-Rabbit IgG (Invitrogen, A-11011), Alexa Fluor® 568 Goat Anti-rat IgG (Invitrogen, A-11077), and Alexa Fluor® 568 Goat Anti-mouse IgG (Invitrogen, A-11004), Alexa Fluor® 488 Donkey anti-sheep IgG, (Invitrogen, A-11015), Alexa Fluor® 546 Donkey anti-Rabbit IgG (Invitrogen, A-10040), Alexa Fluor® 647 Donkey anti-Mouse IgG, (Invitrogen, A-31571).

### Immunohistochemistry

Immunostaining of whole mouse embryos and embryonic sections as well as sex determination via PCR was performed as previously described (Shin et al., 2010; 2014). Concerning the determination of embryo stages, noon of the day when mating plugs were observed was considered as embryonic day 0.5 of development (E0.5).

### Wholemount In situ hybridizations of mouse embryos

In situ hybridizations on dissected embryos were performed as previously described (Henrique et al., 1995; Yoon et al., 2013), including embryo dissections and the preparation and hybridization of probes. Briefly, embryos were dissected at stages E5.5 and E6.5. Noon of the day when mating plugs were observed was considered as embryonic day E0.5. Using forceps, dissections were performed in the dissection medium (DMEM (Gibco) containing 20 mM Hepes, 10% FCS, 100 U/ml penicillin and 100 μg/ml Streptomycin). Established plasmids have been described for making antisense probes recognizing *Wnt-7b* (Yoon et al., 2013), *Mash2, Esrrb* and *Fgf4* (Guzman-Ayala et al., 2004).

## Acknowledgments

We are grateful to K. Tremblay and J. Rossant for sharing plasmids for ISH probes, T. Fazzio, P. Kaufman and E. Torres for advice and discussion. This work was supported from NIH grants R35 GM145263 to I.B., HD083311 to J.M., and G.M.F. was supported by a Wellcome Trust/Royal Society Sir Henry Dale Fellowship (211209/Z/18/Z) and a Medical Research Council New Investigator Award (MR/N000609/1). I.B. is a member of the University of Massachusetts DERC (DK32520).

## Autor contributions

Conceptualization: I.B., F.W.; Investigation, F.W., A.C., Y.Y., M.C.W., C.E-S., F.B.; Supervision: I.B., J.M., and G.M.F; Writing: I.B., F.W.

## Supplemental Figure Legends

**Supplemental Figure 1:**
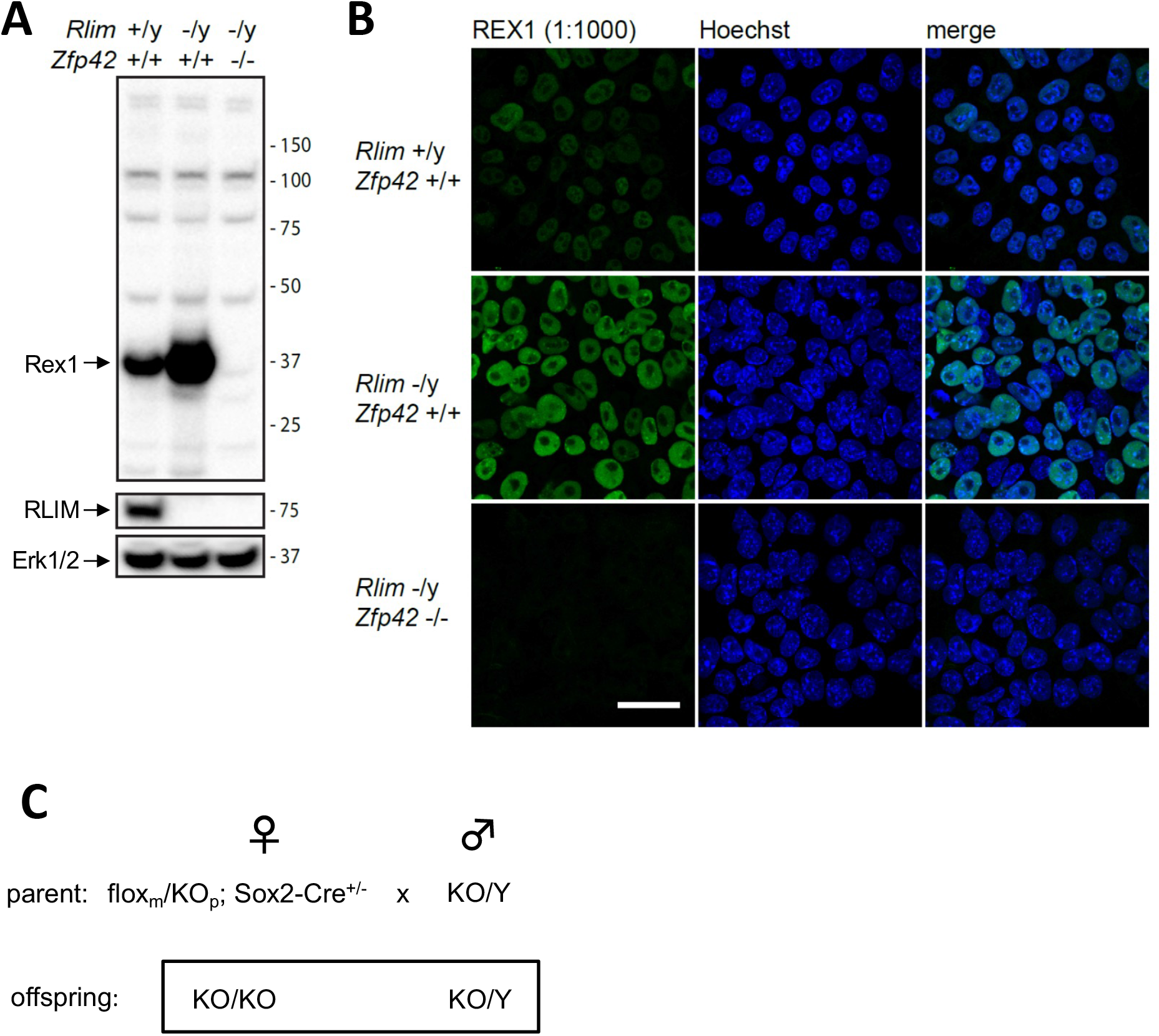
High specificity of Rex1 antibodies. Rex1 antibodies generated in sheep (3^rd^ bleed) were tested using male ESC models including ESCs WT for *Rlim* and Rex1 (*Rlim* +/y; *Zfp42* +/+), ESCs lacking *Rlim* (*Rlim* -/y; *Zfp42* +/+), and ESCs lacking both (*Rlim* -/Y; *Zfp42* -/-) via **A)** Western blotting, and **B)** immunostaining. **C)** Efficient generation of mouse offspring lacking *Rlim*. Parental and offspring genotypes are indicated.

**Supplemental Figure 2:**
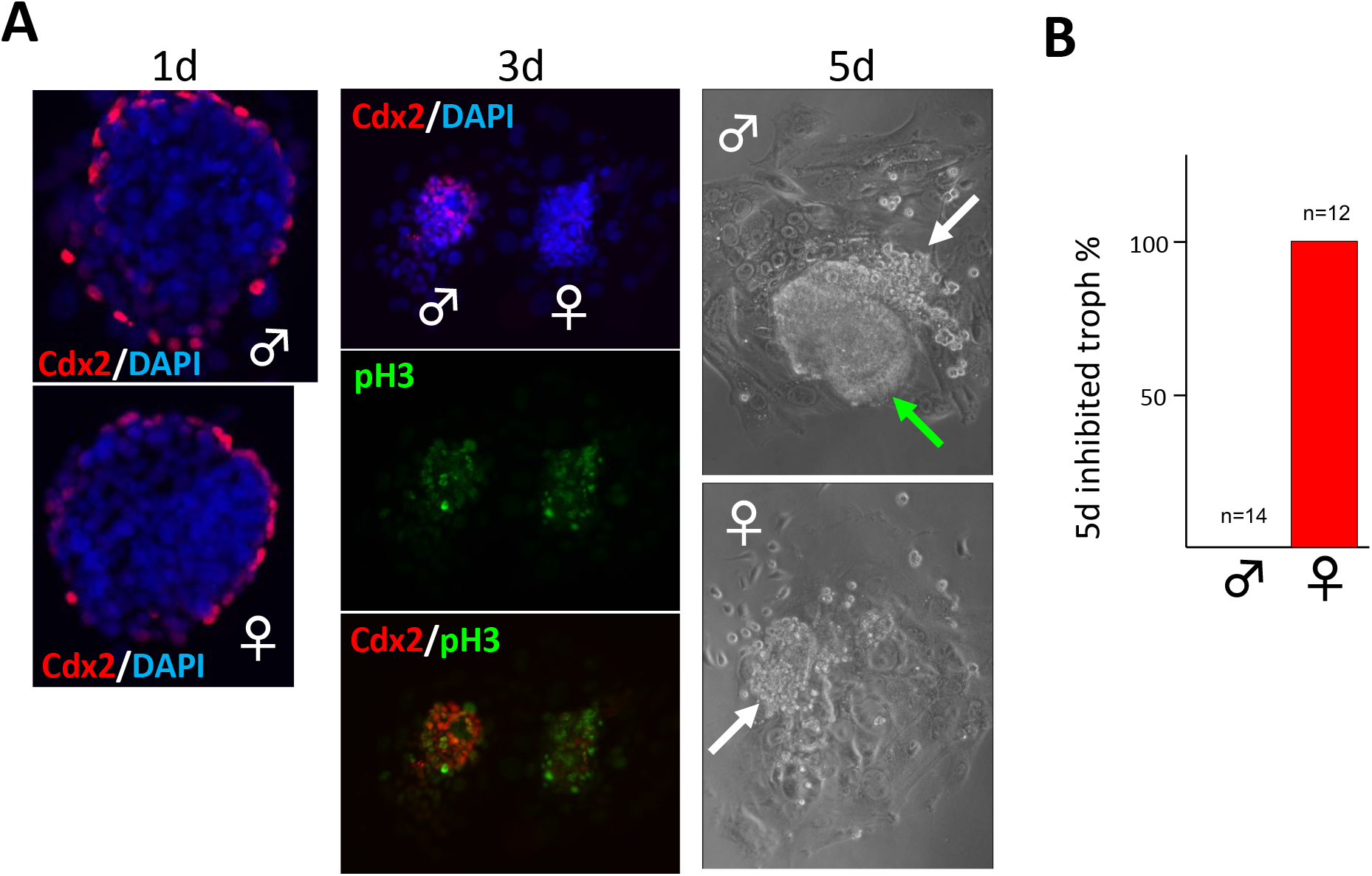
Collapse of Cdx2^high^ cell pool and defective pTE function in female *Rlim*KO blastocyst outgrowths. **A)** Male/female *Rlim*KO blastocysts cultured in vitro. While similar numbers of Cdx2^high^ cells are detected in male and female blastocyst outgrowths cultured for 1 day (**<u>d1)</u>**, by **<u>3d</u>** females start losing Cdx2^high^ trophoblast cells (mostly lost by 4d), with similar numbers of pH3^+^ mitotic cells. At **<u>5d</u>** (brightfield) males but not females have formed large, presumably pTE-derived structures (green arrow). White arrows indicate disorganized epiblast cells. n>10, each. Genotyping after image recording. **B)** Quantification of outgrowths exhibiting inhibited pTE-derived cell growths at 5d.

